# Concerning suppression and facilitation of motion perception in humans

**DOI:** 10.1101/495291

**Authors:** Michael-Paul Schallmo, Scott O. Murray

## Abstract

Here we consider some topics from a commentary by Tzvetanov (2018) on our article entitled “Suppression and facilitation of human neural responses” (Schallmo et al., 2018). The author raises two main points of criticism, which we will summarize thus: 1) psychophysical response bias is not adequately modeled, and 2) other computational models may be better suited to describe the psychophysical motion discrimination data. Both of these points are reasonable in principle, but we disagree with the author in a few areas. We provide some discussion of both issues in order to further elevate the scientific dialogue.

## Introduction to issue #1 - response bias

The issue of modeling response bias within psychophysical data is both interesting and potentially important. To clarify, response bias in the motion direction discrimination paradigm pioneered by Tadin and colleagues (2003) and used in our recent work (Schallmo et al., 2018) would manifest as a tendency for a subject to favor one response direction over the other (i.e., right vs. left) given equal evidence or stimulus information for each. As shown by Tzvetanov (2018) (Figure 2A), this is equivalent to a small shift of the psychometric function along the x-axis. While it is expected that such response biases would be very small for most neuro-typical adults (an x-axis shift of perhaps less than 5 ms), when comparing data between control subjects and individuals from special populations (e.g., schizophrenia), the effect of different response biases between groups may not be trivial.

We also note that the issue of modeling response bias is not limited to the motion discrimination paradigm developed by Tadin and colleagues (2003). Indeed, this issue appears relevant for any 2-alternative forced choice paradigm wherein a bias may be expressed (e.g., choosing the first stimulus in a temporal 2-interval forced choice task). Thus, considering this issue may be of interest to researchers using other psychophysical paradigms.

## Methods and results - examining response bias

Because our experiment was not designed with measuring response bias in mind, doing so post hoc becomes more difficult. Specifically, because our data were acquired using an adaptive staircase procedure with a Weibull function as the assumed form of the underlying psychometric function, there was generally sparse sampling of trials at extremely short stimulus durations (< 20 ms). This is because our chosen adaptive staircase procedure (Psi method, implemented in the Palamedes toolbox; Prins & Kingdom, 2009) will automatically tend to present trials at stimulus durations near the estimated threshold, which in the case of the Weibull function is the duration at which the subject performs with 79% accuracy (Kingdom & Prins, 2010). Further, although the number of trials with left- and right-drifting stimuli was counterbalanced for each stimulus condition in our task, the nature of the adaptive staircase procedure did not permit us to acquire a balanced number of left and right trials for each stimulus duration within a given condition. A low number of trials (< 5) at very short stimulus durations, and unequal sampling of left and right trials, often made it unfeasible to obtain reliable fits to a Logistic model function with threshold values (reflecting the absolute response bias) in the expected range of ± 5 ms. Thus, for many of our data sets we found it was not possible to assess the degree of response bias post hoc. In a subset of data sets which happened to include a sufficient number of short duration trials with a relatively even distribution of left and right drifting stimuli, we observed very small effects of modeling response bias using the method proposed by Tzvetanov (2018) (i.e., changes in threshold estimates on the order of 5 ms; see Figure 1). This agrees with anecdotal observations from Tadin’s laboratory, who found no substantive response biases for foveal motion discrimination among experienced psychophysical observers (Tadin, personal communication).

**Figure 1.**
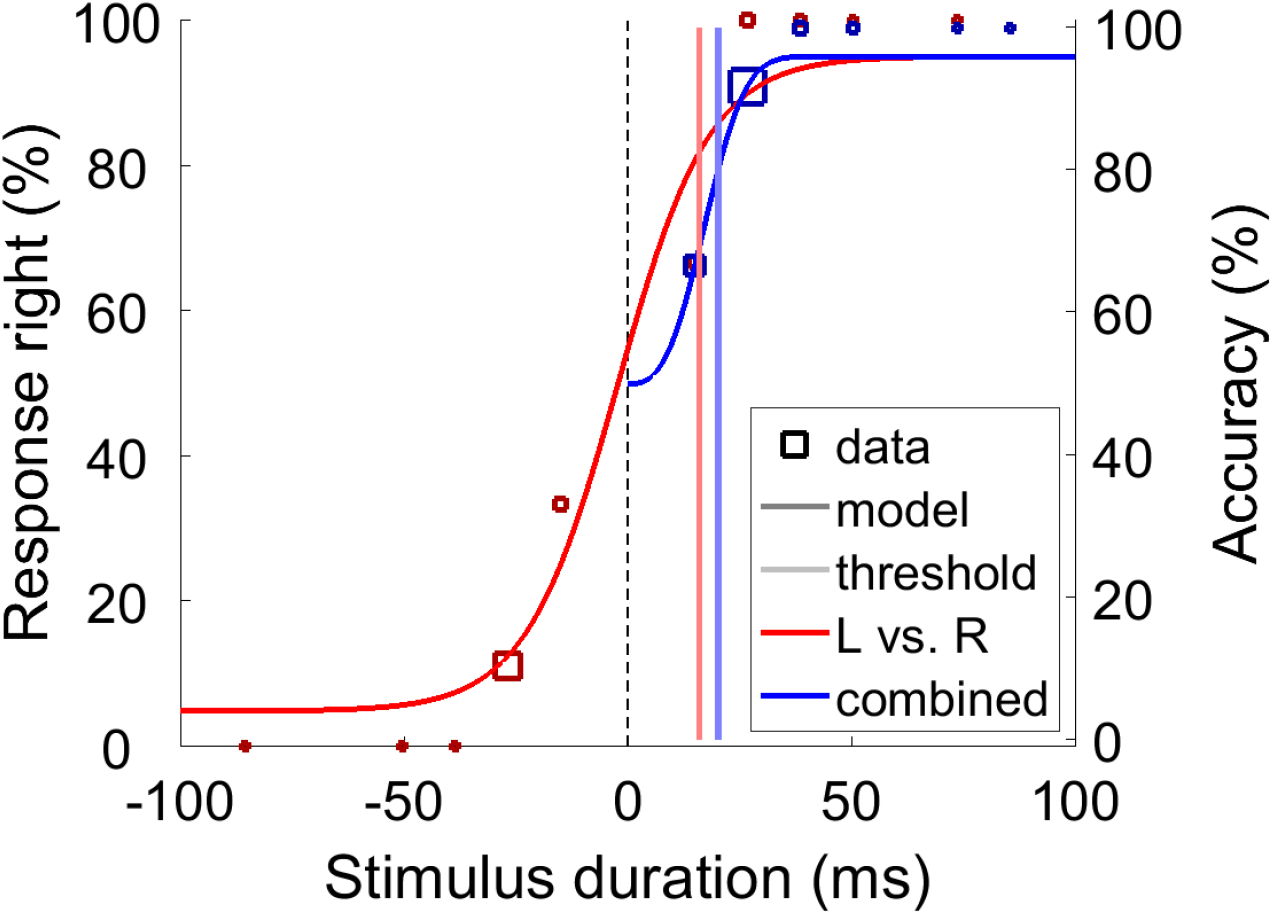
Example data and fits. Psychophysical data are from a single staircase in a single subject in our motion direction discrimination paradigm (Schallmo et al., 2018). Squares show response data (size indicates number of trials), curves show fit psychometric functions, and vertical lines show threshold values (80% accuracy). In red, we show data for left- (negative x) and rightward (positive x) drifting stimuli separately, which were used to model response bias (Tzvetanov, 2018). In blue, we show data for the two directions combined, as in our previous work (Schallmo et al., 2018).

## Discussion - response bias

One issue not addressed by Tzvetanov (2018) is how best to incorporate an estimate of response bias into the psychophysical experiment. This issue bears consideration, as it is easy to imagine a case in which modeling response bias explicitly during data collection might introduce other problems. Although the Logistic model described by Tzvetanov (2018) seems well-suited to a method of constant stimuli approach, using this model within a Psi adaptive staircase procedure (as in our paradigm) would be expected to yield poor estimates of the ‘true’ duration threshold (i.e., the duration at which a subject achieves 80% discrimination accuracy). This is because the Psi method automatically attempts to sample a large number of trials near the threshold, as noted above. In this case, the consequence would be the presentation of a large number of trials at extremely low durations (where accuracy is expected to be near 50%, the threshold value for the Logistic function), and fewer trials at duration values near the 80% accuracy point. Additionally, in the response bias model proposed by Tzvetanov (2018), shifts in the Logistic threshold value reflect changes in response bias only, while differences in the ‘true’ duration threshold (80% accuracy) are captured by changes in the slope parameter. This is another reason for caution regarding the implementation of the response bias Logistic model within a Psi staircase method, as a relatively large number of trials (hundreds) per condition are required to obtain highly reliable slope estimates (Kingdom & Prins, 2010).

We suggest an alternative method for estimating response bias in a motion direction discrimination paradigm within a Psi adaptive staircase framework. By presenting a fixed number of trials (we recommend at least 40 in total) in which grating stimuli are presented for a very short duration (e.g., 10 ms) but do not drift, an estimate of response bias may be obtained. By calculating the ratio of left vs. right responses for these trials in which the stimuli lack directional motion information, one can measure the response bias at ‘drift duration = 0.’ This estimate of response bias could then be combined with data acquired using the Psi method described in our original paper (Schallmo et al., 2018) within the Logistic modeling approach described by Tzvetanov (2018). A similar approach has been used by Zhang, Kwon, and Tadin (2013) to quantify a substantial bias toward perceiving centrifugal motion for static stimuli in the periphery (40° eccentricity). We believe that the approach we suggest above would preserve the advantage of the Psi method to estimate psychometric functions using a relatively small number of trials, while also adequately modeling possible response biases during foveal motion discrimination.

## Introduction to issue #2 - computational modeling

The second criticism raised by Tzvetanov (2018) may be summarized by saying that a different computational framework from the normalization model of Reynolds and Heeger (2009) provides a better description of the direction discrimination psychophysical data. This premise is entirely reasonable, but to some extent it misses the point of themodeling efforts in our original paper. We adopted the Reynolds and Heeger (2009) normalization model in an effort to: 1) present the simplest model that could plausibly account for the spatial suppression phenomena observed psychophysically, 2) determine whether normalization could account for spatial summation (in addition to suppression), and 3) establish a model framework within which we could more easily interpret the results from our experiments using lorazepam and MR spectroscopy, which investigated the role of GABAergic inhibition in the psychophysical spatial suppression effect. Our modeling work builds directly upon that of Rosenberg and colleagues (2015), who showed that weaker divisive normalization could plausibly account for enhanced motion discrimination performance in subjects with autism spectrum disorders, as reported by Foss-Feig and colleagues (2013).

## Discussion - computational modeling

In adopting the normalization model, we acknowledge that it most certainly does not capture the full complexity of how motion discrimination is implemented at a neural level. Divisive normalization is a descriptive computational principle, not a biophysical model of neural circuitry *per se*. Tzvetanov (2018) states that our paper claims that spatial suppression is “not necessarily due to excitatory and inhibitory interactions between motion sensitive neurons but instead arises from the divisive normalization of neuronal activity.” This represents a misunderstanding of our work. It is certainly the case that a series of excitatory and inhibitory neural processes (and likely neuromodulatory functions) give rise to spatial suppression during motion direction discrimination, as these are the foundational elements of the neural circuits underlying this behavior. Our findings are more properly summarized as: 1) the normalization model is sufficient to predict both spatial suppression and summation effects, 2) stronger inhibition, whether driven by pharmacology or by differences in endogenous GABA between individuals, is not associated with stronger spatial suppression (more on this below). Instead, the effects of stronger inhibition may be described within our normalization model framework (Schallmo et al., 2018) in terms of reduced contrast gain (in the case of lorazepam; Figure 4 supplement 1), or a reduced response criterion (in the case of hMT+ GABA levels; Figure 5 supplement 6).

As noted in the Introduction to our original paper (Schallmo et al., 2018), strong assumptions have been made about the relationship between the behavioral phenomenon of spatial suppression and the underlying neural processes (i.e., inhibition). To be clear, previous work has interpreted weaker spatial suppression as evidence for weaker inhibition within particular clinical populations. A primary goal of our original paper was to determine whether there was any evidence to support the idea that inhibition, which is specifically the result of GABAergic neural activity in cortex, is the primary factor that determines the strength of spatial suppression. In failing to find such evidence from our experiments using lorazepam and MR spectroscopy, we do not claim that inhibition is not involved in spatial suppression. Instead, we find that the magnitude of spatial suppression is not a reliable index of the strength of inhibition.

After presenting an alternative computational model to the divisive normalization framework, Tzvetanov (2018) goes on to critique our interpretation of our lorazepam data in light of the newly proposed model results. The author states “duration thresholds of the drug-intake condition increased when compared to the placebo condition. Thus, contrary to their conclusion…their results are consistent with the original hypothesis of drug-based inhibition enhancement of spatial suppression.” This is clearly incorrect. It is apparent from our data (Schallmo et al., 2018) that enhancing GABAergic inhibition via lorazepam had an effect on motion discrimination (see Figure 4). However, when taking lorazepam subjects clearly show weaker spatial suppression in the psychophysical paradigm, which is measured by the difference in thresholds between smaller and larger size conditions (Figure 4C). These data speak for themselves, and any conclusion that lorazepam enhanced spatial suppression is untenable.

Regarding the effort by Tzvetanov (2018) to explicitly model inhibition in terms of specific model parameters, we note that other neural mechanisms of suppression have also been proposed; for example, withdrawal of recurrent excitation as a driving factor behind surround suppression in visual cortex has been clearly modeled and described (Sato, Haider, Haüsser, & Carandini, 2016; Shushruth et al., 2012). Further, there are at least three distinct sub-classes of GABAergic inhibitory neurons (parvalbumin-, somatostatin-, and vasoactive intestinal peptide-positive) that are each thought to play distinct functional roles within visual cortical circuits that exhibit surround suppression (Atallah, Bruns, Carandini, & Scanziani, 2012; Ma et al., 2010; Nienborg et al., 2013). Appreciating this complexity and the difficulty of studying cortical physiology in human subjects *in vivo*, we eschewed an attempt to directly model the role of inhibition in spatial suppression, and instead asked ‘given the normalization model framework, how can the observed effects of inhibition (from the lorazepam and spectroscopy experiments) on motion discrimination performance be described?’ The model presented by Tzvetanov (2018), which follows in the vein of Betts and colleagues (2012), is well-intended, but utilizes too many free parameters for our purposes.

